# The relationship between object-based spatial ability and virtual navigation performance

**DOI:** 10.1101/2023.03.16.532962

**Authors:** Tanya Garg, Pablo Fernández Velasco, Zita Eva Patai, Charlotte P. Malcolm, Victor Kovalets, Veronique D. Bohbot, Antoine Coutrot, Mary Hegarty, Michael Hornberger, Hugo J. Spiers

## Abstract

Spatial navigation is a multi-faceted behaviour drawing on many different aspects of cognition. Visuospatial abilities, such as spatial working memory and mental rotation, in particular, may be key factors. A range of tests have been developed to assess visuospatial processing and memory, but how such tests relate to navigation ability remains unclear. This understanding is important to advance tests of navigation for disease monitoring in Alzheimer’s Disease, where disorientation is an early symptom. Here, we report the use of an established mobile gaming app, Sea Hero Quest, as a measure of navigation ability. We used three separate tests of navigation embedded in the game: wayfinding, path integration and spatial memory in a radial arm maze. In the same participants, we also collected measures of mental rotation (Mental Rotation Test), visuospatial processing (Design Organization Test) and visuospatial working memory (Digital Corsi). We found few strong correlations across our measures. Being good at wayfinding in a virtual navigation test does not mean an individual will rate themself as a confident navigator, be good at path integration, or have a superior memory in a radial arm maze. However, those good at wayfinding tend to perform well on the three visuospatial tasks examined here, and to also use a landmark strategy in the radial maze task. These findings help clarify the inter-relationships between different abilities supporting visuospatial and navigation skills.

**Highlights:**

- Three navigation tests embedded in the game Sea Hero Quest were examined in relation to three object-based visuospatial tasks, and self-ratings of navigation ability and stress during navigation.
- No associations were observed among performance on wayfinding, path integration and radial arm maze levels of Sea Hero Quest.
- Object-based visuospatial abilities were selectively correlated with performance on wayfinding levels of Sea Hero Quest.
- Gameplay stress and navigation strategy were not associated with performance on Sea Hero Quest navigation tasks.

## 1. Introduction

The ability to find our way in different environments is a fundamental skill reliant on multiple cognitive domains. Human navigation involves a host of capacities ranging from planning routes to reading maps, and from identifying landmarks to maintaining a sense of direction. Some people are prone to getting lost in novel environments, while others can easily find their way (Burles & Iaria, 2020; Ekstrom et al., 2018; Wolbers & Hegarty, 2010; Weisberg & Newcombe, 2016; Weisberg et al., 2014). Variation also occurs for other cognitive abilities such as visuospatial working memory, mental rotation and perspective-taking (Allen et al., 1996; Hegarty et al., 2006; Muffato et al., 2022). It is important to understand individual differences in navigation ability that may relate to these competencies because deficits in navigation may constitute an early marker of Alzheimer’s Disease (AD) (Coughlan et al., 2018). There are also negative effects of disorientation and its associated risks (Burles & Iaria, 2020; Fernández Velasco & Casati, 2020). More broadly, such an understanding will advance the domain of spatial cognition at large.

Creating a valid standardised test of navigation is not easy. This is because of the difficulties in achieving the required levels of environmental manipulation and experimental control in standard research settings, which are further compounded by the problem of testing large enough cohorts to account for the wide variations in performance. In recent years, virtual reality (VR) and the widespread touch-screen technology on tablet and mobile devices have opened new possibilities for testing. Our team capitalised on these possibilities by developing a set of tests for navigation ability in the form of the video game app Sea Hero Quest (SHQ) (Spiers et al., 2023). We have employed SHQ to test the navigation ability of 3.9 million people across the world (Coutrot et al., 2018; Walkowiak et al., 2022). SHQ has good test-retest reliability (Coughlan et al., 2020), and has been shown to be predictive of real-world navigational performance (Coutrot et al., 2019). So far, studies using SHQ have found that gender differences in navigation ability for a given country can be partly predicted by gender inequality (Coutrot et al., 2018), that people are better at navigating environments that were topologically similar to the ones in which they grew up (Coutrot et al., 2022a) and that spatial navigation peaks at seven hours of sleep later in life (Coutrot et al., 2022b). It has also been used in the study of AD, to detect sub-optimal navigation performance in pre-clinical AD, to classify spatial impairments in healthy participants at a high risk of AD (Coughlan et al., 2019), and to detect those AD patients most prone to disorientation (Puthusseryppady et al., 2022). Finally, SHQ has also been used to detect wayfinding and path integration (PI) deficits in patients with traumatic brain injury (Seton et al., 2023).

Here, we explore how navigational performance as assessed by SHQ relates to other cognitive abilities and profiles. We test participants on three navigation tasks in SHQ: Wayfinding, PI, and the radial arm maze (RAM) test of spatial memory. Participants were also tested on three measures of visuospatial ability: the Mental Rotation Task Form A (MRT-A; Peters et al., 1995), the Design Organization Test (DOT; Killgore et al., 2005), and the Digital Corsi Block-Tapping Task (D-Corsi; Corsi, 1972). Participants also completed the Santa Barbara Sense of Direction Scale (SBSOD; Hegarty et al., 2002), the Navigation Strategies Questionnaire (NSQ; Brunec et al., 2019), a multiple-choice question about their navigation strategy on RAM levels of SHQ and a questionnaire about perceived stress when navigating in SHQ. Table 1 below shows the twenty-two hypothesised relationships between performance on different tasks. We made predictions based on the literature (e.g., Wolbers & Hegarty, 2010). We predicted that wayfinding would most broadly correlate with other measures due the broader variety of demands in that task.

**Table 1.**
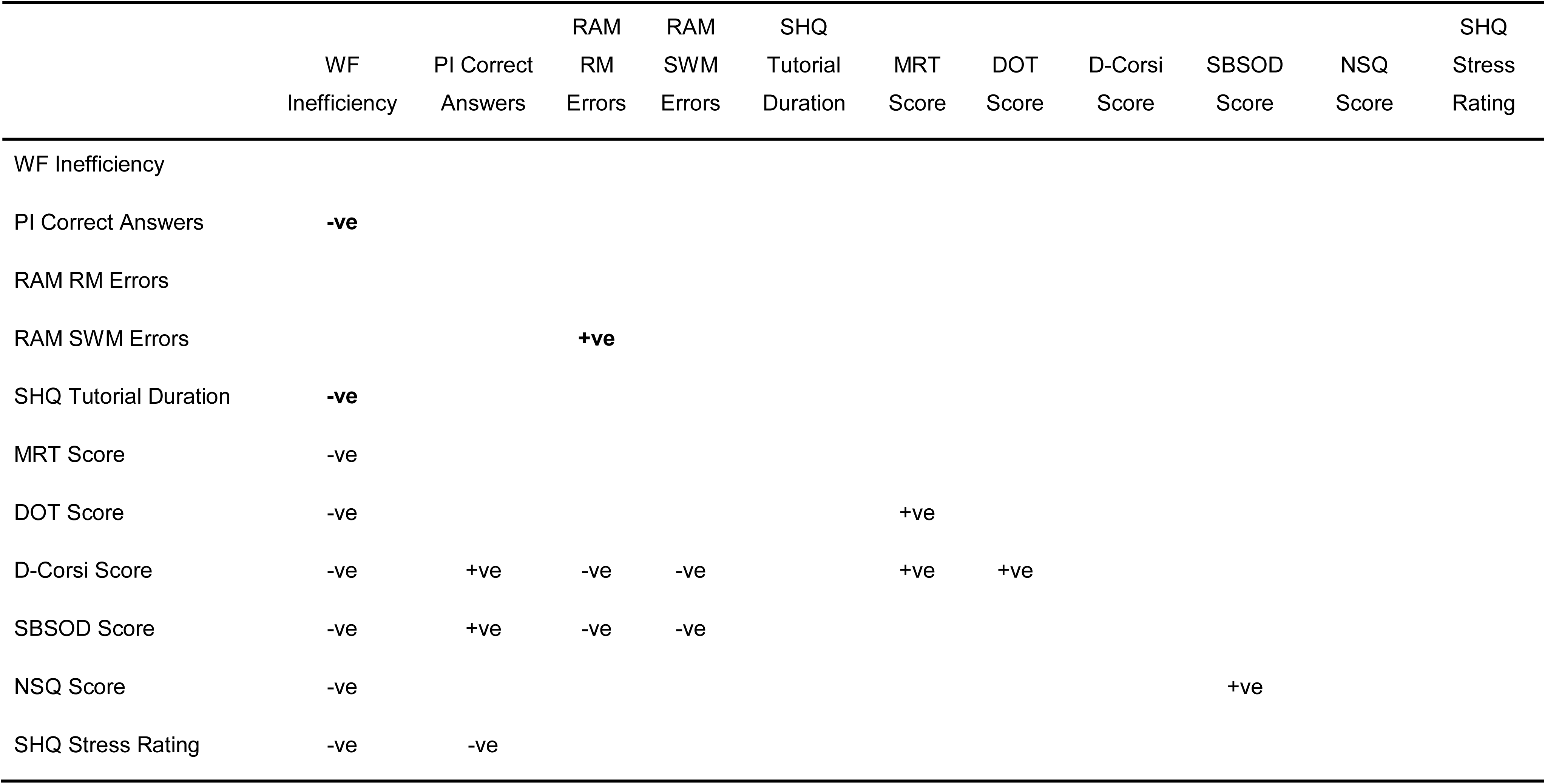
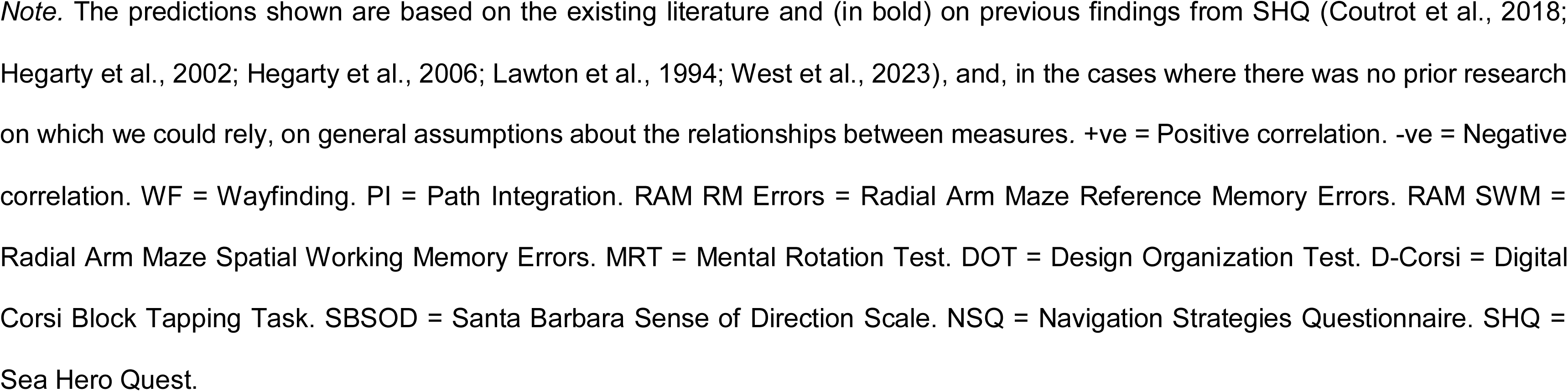
Hypothesised relationships between performance on various tasks.

## 2. Methodology

### 2.1 Context of the Data Collection

This article is an extension of broader research, which focussed on the efficacy of SHQ in reducing the frequency of intrusive memories of analogue trauma using the trauma film paradigm. Reporting of findings relating to SHQ and its effect on intrusive memories is beyond the scope of this paper. The design and analysis of this research were not pre-registered. The data and analysis code for this study will be made publicly available.

### 2.2 Participants

Seventy-eight healthy English-speaking participants over the age of 18 years (*M* = 20.33 years, *SD* = 4.03 years) were recruited from University College London through its online subject pool SONA, and compensated in the form of course credits. Demographic information about the participants is summarised in Table 2. Ethical approval was obtained from the University College London Review Board (9807/004). All participants provided informed written consent.

**Table 2.** Demographics overview for participants in the study.

| Age |  | Gender |  | Race |  |
| --- | --- | --- | --- | --- | --- |
| 18-24 | 73 | Female | 58 | Asian | 45 |
| 25-30 | 4 | Male | 20 | White | 27 |
| 40-49 | 1 |  |  | Other | 4 |
|  |  |  |  | Black | 2 |

### 2.3 Tasks

#### 2.3.1 Sea Hero Quest

##### 2.3.1.1 *Sea Hero Quest Tasks* (Coutrot et al. 2018; Spiers et al., 2023; West et al., 2023)

SHQ is a VR navigation game for mobile and tablet devices in which participants are required to navigate a three-dimensional environment of lakes and rivers. Navigation abilities of participants are assessed through three types of levels: Wayfinding, PI and RAM (see *Figure 1*). Participants played the somewhat “difficult” levels of SHQ. The difficulty of wayfinding levels was based on the number of goals in a particular level, and how far apart they were located from each other. Levels in which there were four or more goals that were located at a considerable distance from each other were considered difficult. For PI levels, the difficulty of levels was determined by the number of turns participants had to take to get from the starting point to the final destination. Levels in which there were at least four turns were selected. Based on the aforementioned criteria, we chose levels 16, 37, 32 and 43 for wayfinding, and levels 44, 49, 54, 59, 64, 69 and 74 for PI. For the purpose of this study, we only analysed levels that were played by all participants in the original trauma study. Accordingly, we considered levels 16 and 37 for wayfinding, and levels 44, 49, 54 and 74 for PI. Lastly, participants played all the RAM levels since there are only five such levels in SHQ

**Figure 1.**
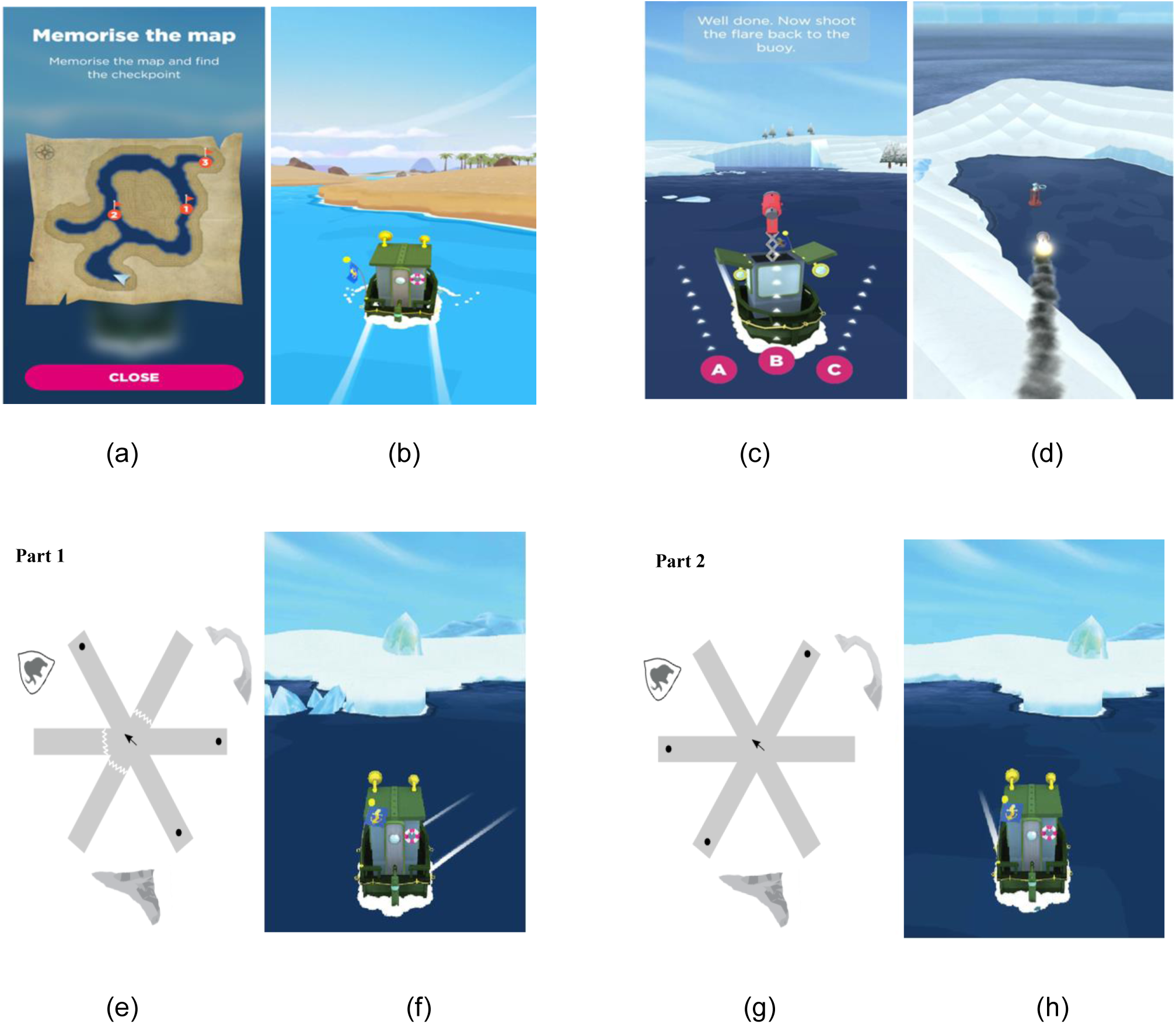
Examples of the various types of levels in Sea Hero Quest. (a-b) Wayfinding Task: Participants memorise a map in the beginning of the task, and navigate to checkpoints in an ordered manner. (c-d): Path Integration (PI) Task: Participants navigate along a river to find a flare gun, and shoot the flare back to the starting point. (e-f) Radial Arm Maze Task (RAM) Part 1: Three of the six arms are blocked, and participants navigate to the three open arms to collect a star that pops out of the water. (g-h): Radial Arm Maze Task Part 2: All six arms are made available, and the participants are required to navigate to the three arms that were blocked during Part 1 to collect the remaining three stars.

In wayfinding levels, participants are required to navigate to checkpoints in an ordered manner based on a map presented to them before the game begins. Performance on wayfinding levels was operationalised as the average inefficiency across all the levels. Lower inefficiency values indicate better wayfinding performance, meaning that participants covered less distance to complete all the levels. To control for prior video gaming experience, we standardised wayfinding inefficiency by dividing it by the duration (in seconds) spent learning SHQ controls in the first two levels of the check-point (practice) task that required no spatial memory to solve. In PI levels, participants navigate along a river to find a flare gun, and shoot the flare back to the starting point by choosing one direction from three alternatives. Performance here was measured as the number of correct answers obtained at the end of all PI levels. Greater number of correct answers indicates better PI performance. The RAM levels are divided into two parts. In Part 1, three of the six arms are blocked, and participants visit the three free arms to collect a star from each of them. In Part 2, the goal of the participants is to visit the three arms that were blocked in Part 1, and collect the remaining three stars from them. Performance on RAM levels was operationalised as the number of reference memory (RM) and spatial working memory (SWM) “errors” made across all the RAM levels. RM errors refer to the number of visits to the arms already visited in Part 1 that needed to be avoided in Part 2. SWM errors refer to the number of visits made to arms in Part 2 that had already been visited during the second part itself. Fewer errors indicate better performance.

##### 2.3.1.2 Sea Hero Quest Stress Rating

Participants were asked to rate the stress they experienced while playing SHQ levels on a scale ranging from 0 (“*not at all”*) to 10 (“*extremely”*) once they completed gameplay.

#### 2.3.2 Visuospatial Ability

The visuospatial abilities of participants were measured using three tasks, which are as follows:

##### 2.3.2.1 Mental Rotation Test-A (Peters et al., 1995)

The MRT-A consists of 12 stimuli, each of which is a two-dimensional image of a three-dimensional object drawn by a computer. We modified the MRT-A to include three answer options and only one correct answer per question instead of four answer options and two correct answers, respectively (see *Figure 2*). We also reduced the time limit from 4 minutes to 3 minutes to maintain adequate time pressure. Participants selected the answer option they thought was the rotated version of the target stimulus. Scores range from 0 to 12. Higher scores indicate better performance.

**Figure 2.**
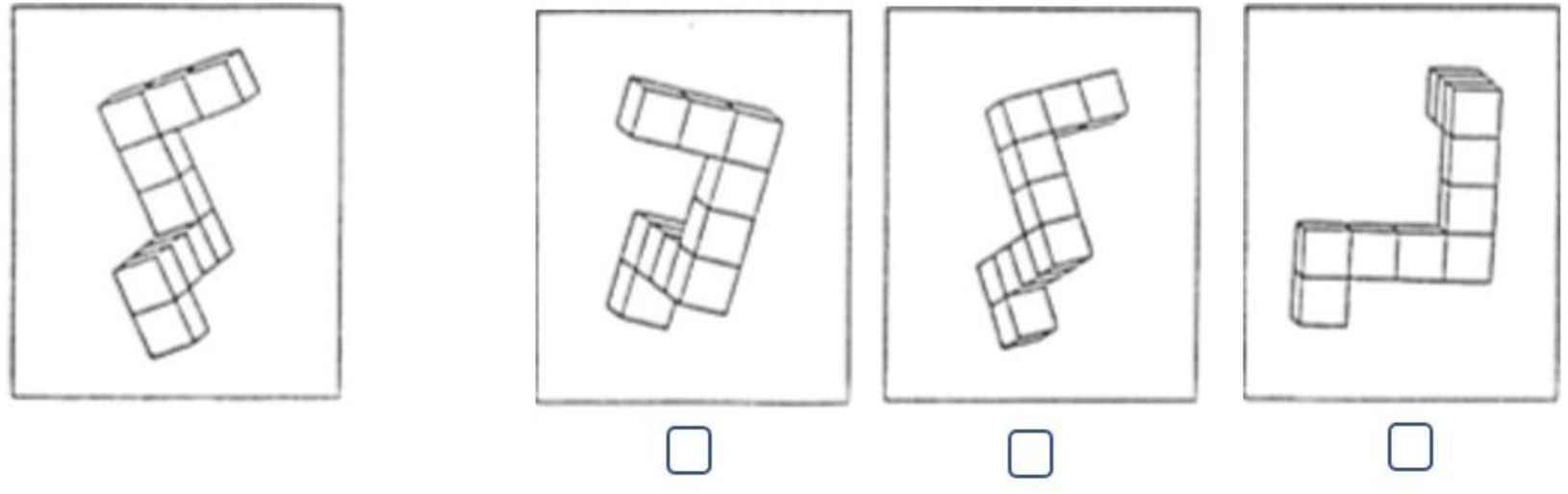
Example item from the modified Mental Rotation Test Form A.

##### 2.3.2.2 Design Organization Test (Killgore, Glahn & Casasanto, 2005)

Participants completed both Form A and Form B of the DOT in a counterbalanced manner. At the top of the page, there is a row of six squares that contains a numerical code key from 1 to 6. There are nine square grids below the code key, each of which showcases a unique pattern that consists of a specific combination of the six squares in the code key (see *Figure 3*). Below each grid was an identically-sized empty grid, which participants completed using numbers from the code key that corresponded to the pattern above. As it is possible to get a ceiling effect within two minutes, participants were allotted one minute to complete this measure (Burggraaf et al., 2016). Scores range from 0 to 112. Higher scores indicate better performance.

**Figure 3.**
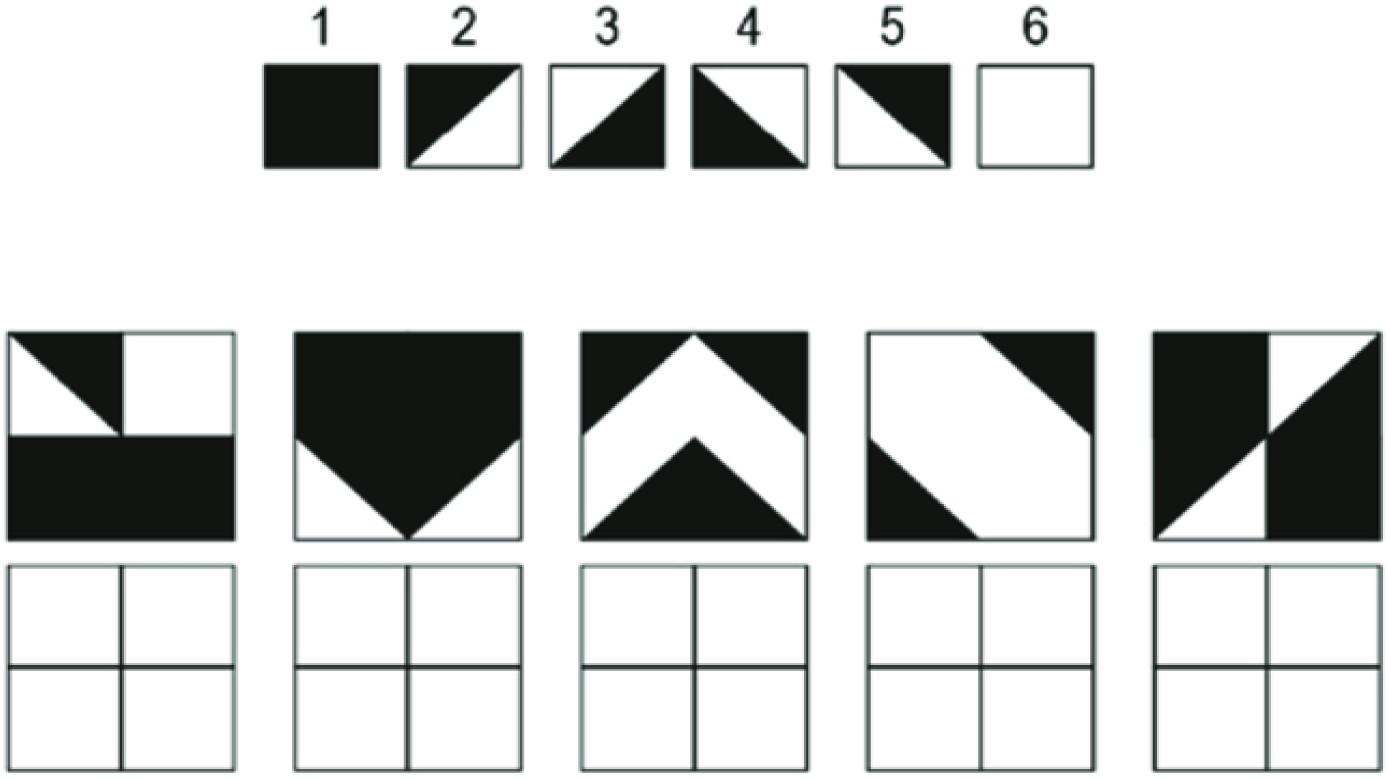
Code key and practice items from the Design Organization Test Form A. From William D. S. Killgore, David C. Glahn & Daniel J. Casasanto (2005). Development and Validation of the Design Organization Test (DOT): A Rapid Screening Instrument for Assessing Visuospatial Ability, *Journal of Clinical and Experimental Neuropsychology*, 27:4, 449-459, https://doi.org/10.1080/13803390490520436. Copyright 2006 by Taylor & Francis Ltd (www.tandfonline.com). Reprinted with permission.

##### 2.3.2.3 Digital Corsi (Corsi, 1972)

In this task, participants observed a set of nine blocks on the computer screen. A number of the observed blocks lit up in a particular sequence, starting with three blocks and increasing with every successful trial (see *Figure 4*). Participants then repeated the sequence by clicking on the blocks in the order in which they lit up. The sum of the number of blocks they clicked in the correct order was computed as their total score, which could range from 0 to 150. Higher scores indicate better performance.

**Figure 4.**
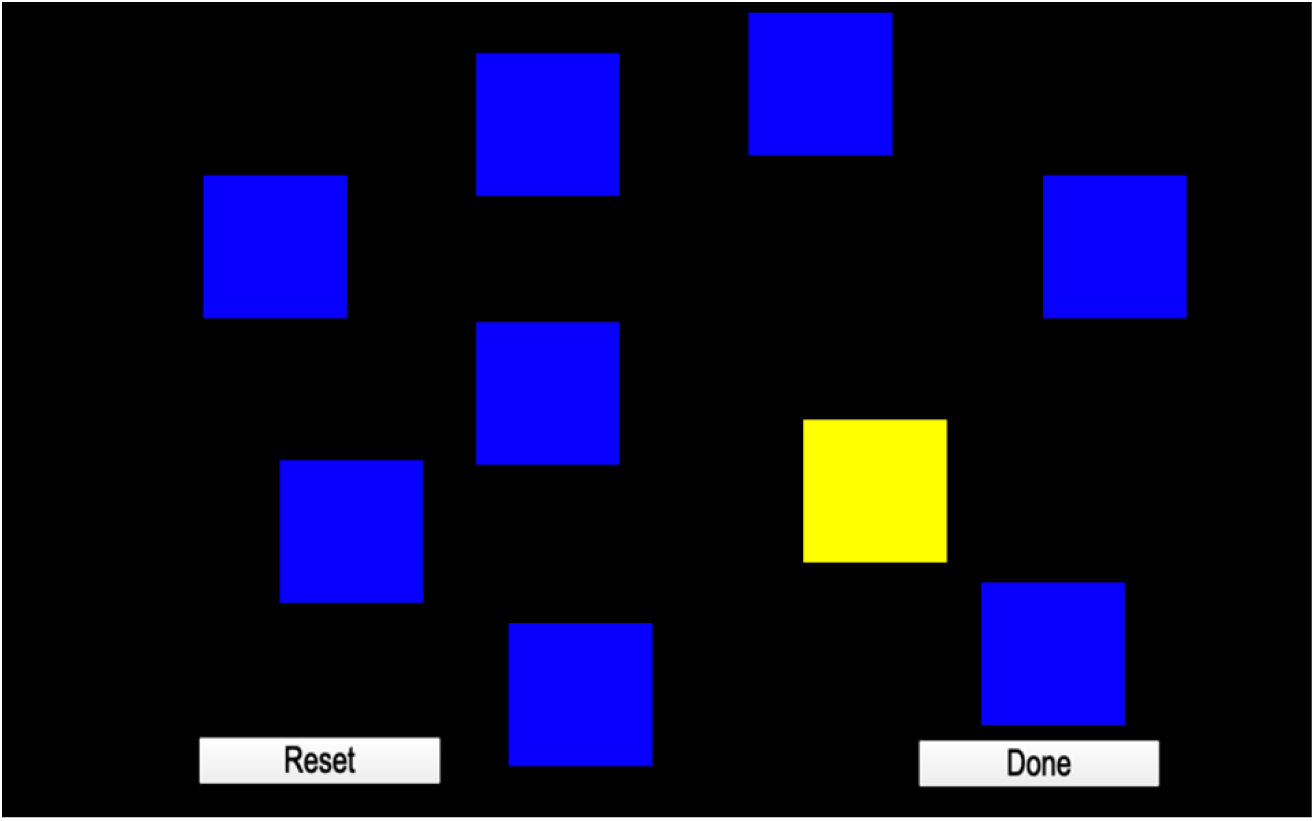
Example of a block lighting up in the Digital Corsi Task. Material extracted and modulated from INQUISIT software package 6.0 (Millisecond Software, LLC, Seattle, WA, United States) with permission from the owner.

#### 2.3.3 Navigation-related Information

Information about navigation preferences and strategies of participants was obtained using the measures listed below.

##### 2.3.3.1 Santa Barbara Sense of Direction Scale (Hegarty et al., 2002)

The SBSOD was used to assess the perceived sense of direction of participants. The instrument consists of 15 items, which were measured on a seven-point scale (“*strongly agree*” to “*strongly disagree*”). Total scores range from 1 to 7. Higher scores indicate better perceived sense of direction. The SBSOD has been demonstrated to have high internal consistency (α = .88) and test-retest reliability (α = .91), which was assessed by administering the questionnaire 40 days apart. Construct validity was determined by significant correlations between SBSOD scores and wayfinding performance.

##### 2.3.3.2 Navigation Strategies Questionnaire (Brunec et al., 2019)

The NSQ consists of 14 items that evaluate the propensity for map-based navigation. The difference between the number of map-based answers and non-map-based answers is taken as a measure of mapping tendency. Total scores range from -14 to +14. Higher scores represent greater use of map-based strategy during navigation.

##### 2.3.3.3 Radial Arm Maze Navigation Strategy

After the completion of every RAM level, participants were presented with a multiple-choice question, which required them to indicate the type of navigation strategy they used to complete the level. Participants were asked, “*How did you navigate? (1) Counted from the start; (2) Used multiple landmarks; (3) Counted from the landmark*.*”* We recorded the most frequently used navigation strategy. Participants were categorised as having used a landmark strategy if they indicated either of the latter two options on at least three out of f ive RAM levels. The same was done for counting strategy if the first option was indicated on the majority of RAM levels. If participants skipped a question, and their dominant navigation strategy could not be determined (i.e., they had used counting strategy and landmark strategy on two levels each), their data were excluded from the analysis.

### 2.4. Procedure

The study was conducted over two sessions, which were held a week apart from each other. Before the first session, participants attempted the modified MRT-A to provide an initial measure of their visuospatial ability. In the first experimental session, participants played SHQ wayfinding and PI levels for six minutes each. Before starting SHQ gameplay, they completed practice levels as part of a brief tutorial to learn the controls of the game. At the end of the session, participants indicated their stress level during gameplay, and attempted the DOT. Upon their return a week later, participants played RAM levels, and indicated their navigation strategy for each level. They also performed the D-Corsi, and completed the SBSOD and the NSQ.

### 2.5 Power Analysis

Based on the estimates from the prior work motivating the approach, the present study assumed the effect size of *d* = .80 (James et al. 2015). Using the conversion calculations by Ruscio (2008), we determined that *d* = .80 amounts to *r* = 0.371. Sample size calculations using these values revealed that a minimum of 72 participants were required to achieve 90% power in order to detect a difference at the 5% significance level.

### 2.6 Data Analysis

The data were analysed using IBM SPSS Statistics (Version 28). Extreme outliers that were three or more standard deviations away from the mean of the task performed were excluded from the analysis of the task in question, though their data for other tasks were retained if they were not outliers on them. Pearson correlation analysis was used to explore the relationship between SHQ and all the measures of visuospatial abilities, navigation and stress. Correlations for predicted relationships were evaluated at the *p* < .05 threshold (see Table 1), while all the other correlational analyses were Bonferroni-corrected for multiple comparisons, *p* <.0009 (55 comparisons) to control for Type I error. Independent samples *t*-tests were also performed to examine how performance between those who used landmark strategy or counting strategy most frequently on RAM levels differ on other measures. Finally, Pearson chi-square test was used to examine the relationship between the propensity for map-based navigation (as indicated by the NSQ) and the type of navigation strategy used on RAM levels (used landmark strategy or counting strategy).

## 3. Results

We calculated the scores of participants across all the measures on which they were tested. The descriptive statistics of participant performance are summarised in Table 3 and Table 4. Data from one participant was not recorded due to a technical error.

**Table 3.** Descriptive statistics of participant performance across all measures.

| Measure | Parameter | Overall |  |  | Females |  |  | Males |  |  |
| --- | --- | --- | --- | --- | --- | --- | --- | --- | --- | --- |
|  |  | *n* | *M* | *SD* | *n* | *M* | *SD* | *n* | *M* | *SD* |
| SHQ | WF Inefficiency | 71 | 26.97 | 6.92 | 52 | 27.18 | 7.09 | 19 | 26.41 | 6.61 |
|  | PI Correct Answers | 75 | 1.72 | 0.99 | 55 | 1.71 | 1.01 | 20 | 1.75 | .96 |
|  | RAM RM Errors | 77 | 5.01 | 2.58 | 57 | 5.34 | 2.51 | 20 | 4.07 | 2.61 |
|  | RAM SWM Errors | 77 | .94 | 1.12 | 57 | 1.07 | 1.18 | 20 | .57 | .84 |
|  | SHQ Tutorial Duration | 74 | 46.52 | 6.45 | 55 | 47.36 | 6.81 | 19 | 44.07 | 4.61 |
| VSA | MRT Score | 78 | 7.37 | 3.10 | 58 | 7.07 | 3.24 | 20 | 8.25 | 2.51 |
|  | DOT Score | 76 | 62.21 | 9.06 | 57 | 62.35 | 8.83 | 19 | 61.79 | 9.96 |
|  | D-Corsi Score | 77 | 61.86 | 18.82 | 58 | 60.50 | 17.55 | 19 | 66.00 | 22.26 |
| Navigation Questionnaires | SBSOD Score | 77 | 4.09 | 0.77 | 57 | 4.02 | .74 | 20 | 4.31 | .83 |
|  | NSQ Score | 77 | -1.83 | 5.03 | 57 | -2.44 | 4.80 | 20 | -.10 | 5.37 |
| Gameplay Stress | SHQ Stress Rating | 77 | 3.96 | 2.33 | 57 | 4.44 | 2.39 | 20 | 2.60 | 1.50 |

*Note.* Sex differences are reported in the Appendix.

**Table 4.**
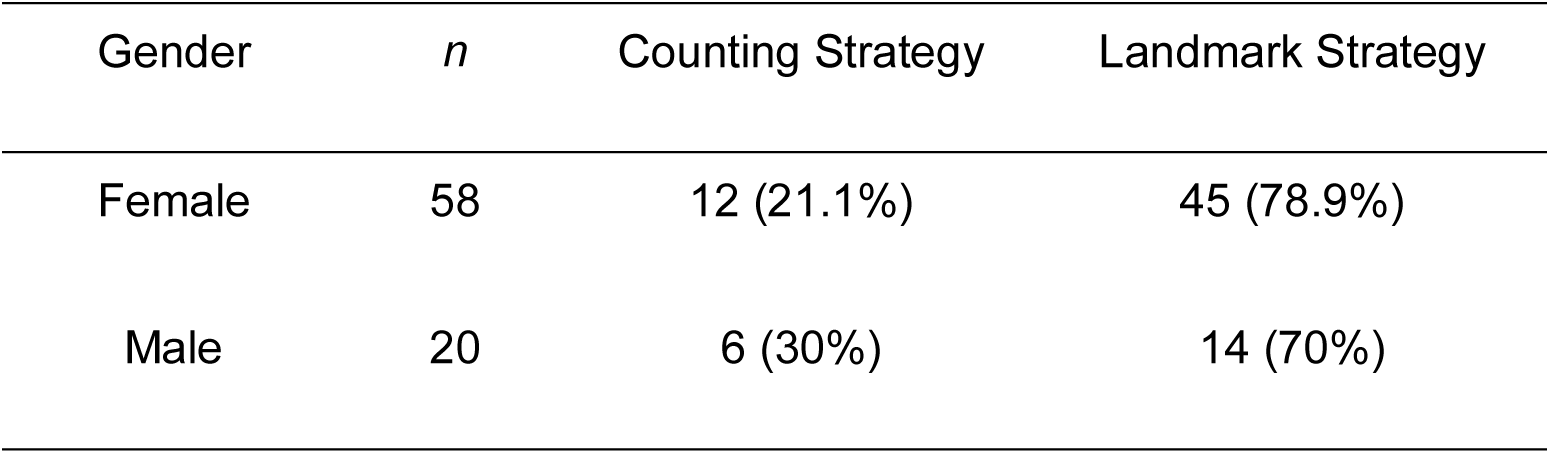
Strategy employed to navigate RAM levels.

| Gender | *n* | Counting Strategy | Landmark Strategy |
| --- | --- | --- | --- |
| Female | 58 | 12 (21.1%) | 45 (78.9%) |
| Male | 20 | 6 (30%) | 14 (70%) |

Pearson correlation analysis revealed that all the measures of visuospatial abilities were associated with each other (*p* < .05, See Table 5). Similarly, responses on the two self-report measures of navigation were related, (*p* < .05, see Table 5). However, visuospatial and navigation measures were not correlated with each other (*p* > .05, See Table 5). Further, visuospatial abilities were only selectively associated with performance on SHQ wayfinding inefficiency. While all visuospatial abilities measures were associated with wayfinding inefficiency, they were not associated with SHQ tutorial duration. Only D-Corsi scores were associated with performance on PI levels. Neither measure of navigation was associated with SHQ wayfinding inefficiency. The significance of the association between navigation and stress did not hold at the Bonferroni-corrected threshold (*p* > .0009). Gameplay stress was also found to be correlated with DOT, but again, the association was not significant at the Bonferroni-corrected threshold (*p* > .0009). The findings from correlation analyses are summarised in Table 5 and *Figure 5*. See the Appendix for further supporting analysis of the large data set reported in Coutrot et al. (2022a). This shows the pattern of correlation between the training levels, PI levels and wayfinding performance varies across levels used in SHQ.

**Figure 5.**
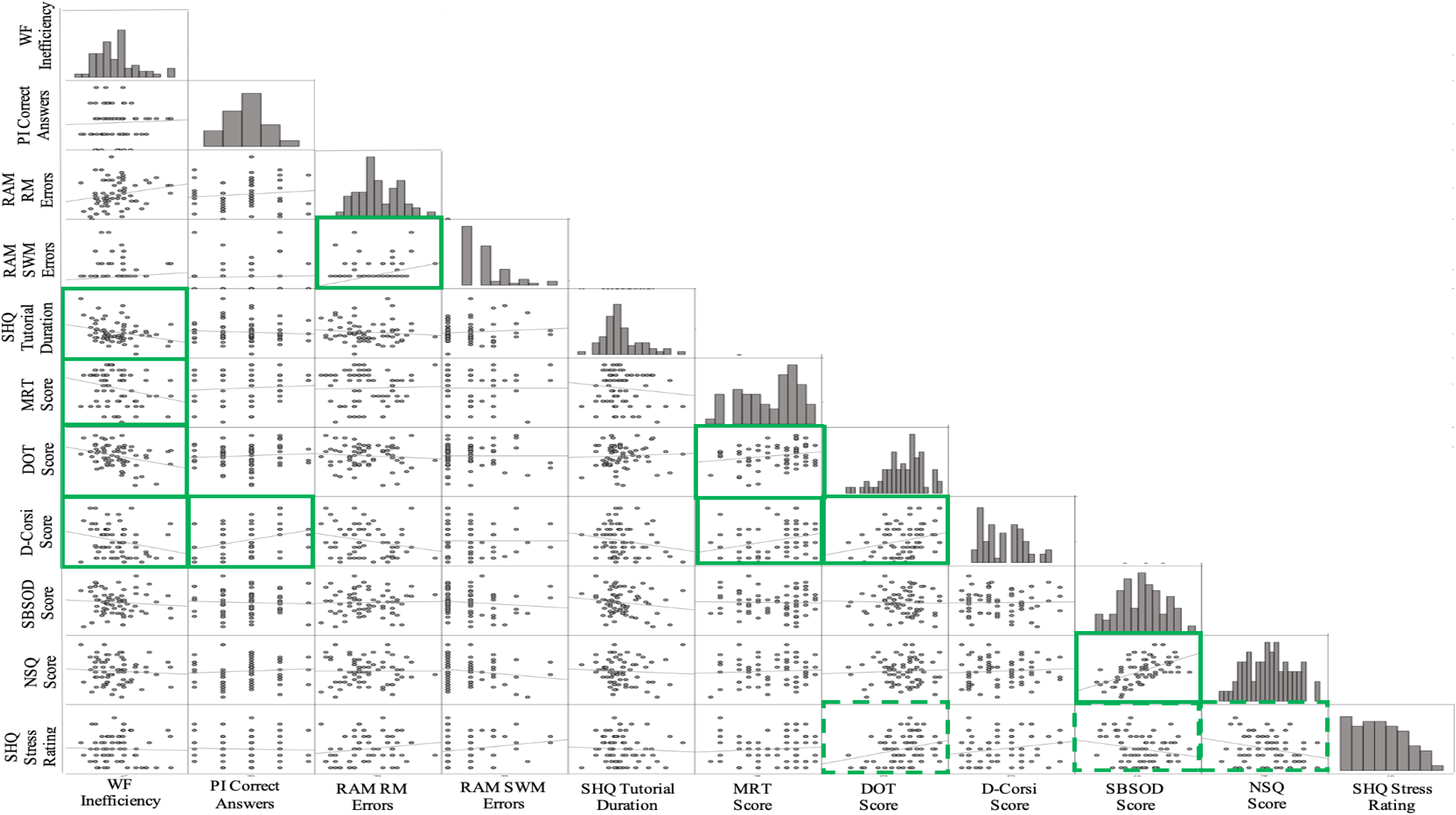
Scatterplot matrix of relationships between 11 measures. Solid green boxes indicate predicted significant relationships. Dotted green boxes indicate unpredicted relationships that were significant at *p* < .05, but did not meet the Bonferroni-corrected threshold for significance (*p* <.0009). WF = Wayfinding. PI = Path Integration. RAM RM Errors = Radial Arm Maze Reference Memory Errors. RAM SWM = Radial Arm Maze Spatial Working Memory Errors. MRT = Mental Rotation Test. DOT = Design Organization Test. D-Corsi = Digital Corsi Block Tapping Task. SBSOD = Santa Barbara Sense of Direction Scale. NSQ = Navigation Strategies Questionnaire. SHQ = Sea Hero Quest. See Appendix for further analysis.

**Table 5.** Correlation matrix for all participants.

|  | WF<br>Inefficiency | PI Correct<br>Answers | RAM RM<br>Errors | RAM SWM<br>Errors | SHQ Tutorial<br>Duration | MRT Score | DOT Score | D-Corsi Score | SBSOD Score | NSQ Score |
| --- | --- | --- | --- | --- | --- | --- | --- | --- | --- | --- |
| PI Correct Answers | 0.053 ($p=.664$) | | | | | | | | | |
| RAM RM Errors | 0.201 ($p=.093$) | 0.142 ($p=.224$) | | | | | | | | |
| RAM SWM Errors | 0.102 ($p=.398$) | -0.012 ($p=.920$) | 0.275 ($p=.016$) | | | | | | | |
| SHQ Tutorial Duration | -0.282 ($p=.018$) | -0.119 ($p=.320$) | -0.143 ($p=.223$) | 0.143 ($p=.223$) | | | | | | |
| MRT Score | -0.267 ($p=.024$) | 0.085 ($p=.470$) | -0.006 ($p=.956$) | 0.001 ($p=.990$) | -0.124 ($p=.292$) | | | | | |
| DOT Score | -0.304 ($p=.010$) | 0.124 ($p=.296$) | -0.060 ($p=.612$) | -0.013 ($p=.914$) | 0.016 ($p=.894$) | 0.269 ($p=.019$) | | | | |
| D-Corsi Score | -0.261 ($p=.029$) | 0.328 ($p=.004$) | -0.199 ($p=.085$) | -0.010 ($p=.930$) | -0.159 ($p=.178$) | 0.301 ($p=.008$) | 0.259 ($p=.025$) | | | |
| SBSOD Score | -0.053 ($p=.658$) | -0.060 ($p=.609$) | -0.024 ($p=.835$) | -0.040 ($p=.732$) | -0.160 ($p=.176$) | 0.039 ($p=.736$) | -0.023 ($p=.844$) | -0.033 ($p=.780$) | | |
| NSQ Score | -0.041 ($p=.734$) | 0.099 ($p=.400$) | -0.048 ($p=.682$) | -0.151 ($p=.193$) | -0.097 ($p=.416$) | 0.100 ($p=.387$) | 0.048 ($p=.683$) | 0.022 ($p=.851$) | 0.577 ($p=.000$) | |
| SHQ Stress Rating | -0.030 ($p=.804$) | -0.017 ($p=.885$) | 0.184 ($p=.110$) | 0.157 ($p=.173$) | -0.049 ($p=.679$) | 0.004 ($p=.972$) | 0.243 ($p=.036$) | 0.142 ($p=.221$) | -0.291 ($p=.011$) | -0.295 ($p=.010$) |

*Note.* The colour intensity of the cells indicates the strength of a correlation, red = negative associations and green = positive. Solid border indicates predicted relationships meeting criteria for significance ( $p < .05$ ). Dotted border indicates unpredicted correlations that were significant at $p < .05$ , but did not meet the Bonferroni-corrected threshold for significance ( $p < .0009$ ). WF = Wayfinding. PI = Path Integration. RAM RM Errors = Radial Arm Maze Reference Memory Errors. RAM SWM = Radial Arm Maze Spatial Working Memory Errors. MRT = Mental Rotation Test. DOT = Design Organization Test. D-Corsi = Digital Corsi Block Tapping Task. SBSOD = Santa Barbara Sense of Direction Scale. NSQ = Navigation Strategies Questionnaire. SHQ = Sea Hero Quest.

Independent 2-tailed *t*-tests revealed that participants who used landmark strategy on RAM levels made significantly fewer RM errors than those who used counting strategy. Such participants also had significantly lower wayfinding inefficiency and higher D-Corsi score on average than participants who employed counting strategy. Table 6 summarises statistics for variables that were significant.

**Table 6.** Metrics found to be significantly different for landmark strategy (n = 59) versus counting strategy (n = 18) on RAM levels. Non-significant metrics not shown.

| Measure | Landmark Strategy |  | Counting Strategy |  | Statistics |  |  |  |
| --- | --- | --- | --- | --- | --- | --- | --- | --- |
|  | *M* | *SD* | *M* | *SD* | *t* | *df* | *p* | *d* |
| WF Inefficiency | 25.71 | 6.50 | 31.00 | 6.88 | 2.88 | 69 | .005 | .802 |
| RAM RM Errors | 4.52 | 2.60 | 6.61 | 1.80 | 3.16 | 75 | .002 | .853 |
| D-Corsi Score | 64.32 | 19.66 | 54.12 | 13.51 | -2.00 | 74 | .049 | -.551 |

Based on the results of Table 6, which are consistent with previous findings of West et al. 2023, additional correlational analyses were conducted to check the association between both types of RAM errors and all SHQ and neuropsychological measures for RAM landmark strategy users only. We found a significant positive association between RAM RM errors and RAM SWM errors. Gameplay stress was also found to be correlated with RAM RM errors, but the association was not significant at the Bonferroni-corrected threshold (*p* > .0009). The findings are summarised in Table 7.

**Table 7.** Correlations for RAM errors for RAM landmark strategy users only.

|  | RAM RM Errors | RAM SWM Errors |
| --- | --- | --- |
| RAM SWM Errors | .352 ($p = .006$) | |
| WF Inefficiency | .131 ($p = .347$) | .117 ($p = .400$) |
| PI Correct Answers | .085 ($p = .529$) | .035 ($p = .794$) |
| SHQ Tutorial Duration | -.070 ($p = .609$) | .112 ($p = .412$) |
| MRT Score | .012 ($p = .930$) | .087 ($p = .511$) |
| DOT Score | -.047 ($p = .725$) | -.040 ($p = .767$) |
| D-Corsi Score | -.131 ($p = .323$) | .034 ($p = .801$) |
| SBSOD Score | -.069 ($p = .609$) | -.064 ($p = .631$) |
| NSQ Score | -.130 ($p = .329$) | -.155 ($p = .246$) |
| SHQ Stress Rating | .324 ($p = .012$) | .205 ($p = .120$) |

*Note.* The colour intensity of the cells indicates the strength of a correlation, red = negative associations and green = positive. Solid border indicates predicted relationships meeting criteria for significance ( $p < .05$ ). Dotted border indicates unpredicted correlations that were significant at $p < .05$ , but did not meet the Bonferroni-corrected threshold for significance ( $p < .0009$ ). WF = Wayfinding. PI = Path Integration. RAM RM Errors = Radial Arm Maze Reference Memory Errors. RAM SWM = Radial Arm Maze Spatial Working Memory Errors. MRT = Mental Rotation Test. DOT = Design Organization Test. D-Corsi = Digital Corsi Block Tapping Task. SBSOD = Santa Barbara Sense of Direction Scale. NSQ = Navigation Strategies Questionnaire. SHQ = Sea Hero Quest.

We did not find an association between mapping tendency as indicated by the NSQ and the navigation strategy used on RAM levels, χ2 (1, N = 77) = .051, *p* = .821.

## 4. Discussion

It would seem obvious that if someone is good at finding their way in a new environment, they should also be good at pointing back to where they started a journey, remembering where objects are in a maze, and rate themselves as a good navigator. Here, we find this is not the case. Rather we find few correlations between different tests of navigation, self-ratings and 2D visuospatial tasks. We tested participants with virtual navigation tasks (wayfinding, PI and RAM) in the app SHQ, visuospatial tasks (MRT, DOT and D-Corsi), and questionnaires about navigation experience (SBSOD, NSQ and RAM navigation strategy) to better understand how these measures may relate to each other. There were weak correlations across the different measures, with little correlation between the three spatial navigation tasks used, and weak correlations between these and self-ratings and navigation strategies. We observed modest correlations between each of the three visuospatial tasks, and all of them with the wayfinding task, but not with other navigation tasks. We discuss what these results mean for understanding cognitive profiles of navigation ability.

Consistent with our predictions, we found a significant correlation between wayfinding performance and performance on visuospatial tasks: MRT, DOT and D-Corsi. To our knowledge, this is the first study to explore the relationship between DOT and navigation tasks. In all three cases, a low-to-moderate correlation is in line with previous studies examining the relationship between large-scale and small-scale spatial abilities (Hegarty et al., 2006; Hegarty & Waller, 2005). The absence of strong correlations between these tests indicates that while wayfinding and these tasks may have some overlap in the cognition required, they also make different demands. This is a pattern discussed in past research exploring the relation between small-scale spatial abilities and large-scale wayfinding ability (Allen et al., 1996; Hegarty et al., 2006). These findings are useful in highlighting the potential utility of VR-based navigation tasks in capturing something that is distinct from object-based visuospatial tasks. One factor that may differentiate wayfinding from the other visuospatial tasks is the reliance on broader executive functions demands. Unlike object-based visuospatial tasks, wayfinding places specific demands on planning and inhibition (Miyake et al., 2001; Patai & Spiers, 2021). In SHQ this involves avoiding re-approaching visited checkpoints and planning optimal paths given the order of checkpoints indicated on the initially shown map.

We had predicted that RAM levels and D-Corsi performance would be correlated because both require holding a set of locations in mind over many seconds i.e., they test spatial working memory. We found no evidence for this. This result is consistent with the view that ‘spatial working memory’ is not a unitary cognitive function, but depends on the context (e.g., 2D screen space versus VR rendered 3D environment). Theoretically, it seems plausible that the D-Corsi is supported mainly via frontoparietal circuits, and the RAM additionally draws on hippocampal circuits to support the representation of the large-scale environment (Iaria et al., 2003; West et al., 2023). The absence of a correlation between the RAM and other navigation tasks further indicates that different cognitive demands between the tasks are involved.

A surprising result is that neither the SBSOD nor the NSQ is correlated with any of the SHQ performance measures. This stands in contrast with previous evidence of correlations between navigation behaviour and these measures (Brunec et al., 2019; Hegarty et al., 2002; Sholl et al., 2006), but is similar to some other evidence (Gerb et al., 2022). It may be that participants need to physically move or use body-based cues to navigate in a space for a stronger association of navigation behaviours with standardised measures such as the SBSOD (e.g., Hegarty et al., 2006). Another possibility is that people tend to rate their navigation ability in relation to how often they might get lost. In the UK sample we tested, it is likely many participants would use GPS-based systems to find their way and might rarely find themselves in the situation simulated in the wayfinding task; where a map is studied and must be committed to memory before navigation. When we recently sampled a large population, drawn from many nations and a range of ages, we found a consistent relationship between wayfinding performance in SHQ and self-rated navigation ability (Walkowiak et al., 2022). Thus, the relationship between self-ratings and wayfinding performance is likely moderated by a wide range of factors (e.g., He & Hegarty, 2020; Hegarty et al., 2006; van der Ham & Koutzmpi, 2022; Van der Ham et al., 2021). Notably, while actual performance in the wayfinding task in SHQ was not correlated with the SBSOD or the NSQ, the stress ratings of playing SHQ were. Thus, people who think they are good navigators, or those who tend to think using maps, will be less likely to find the SHQ tasks stressful. In future, it will be useful to explore more specific questionnaires about spatial anxiety in daily life and SHQ performance to understand if these are linked more than the stress reported (Lawton, 1994; Lawton & Kallai, 2002).

We found that the SBSOD and the NSQ are moderately correlated with each other, which indicates that a self-rating as having a strong sense of direction is associated with reporting using map-based strategy to navigate. To the authors’ knowledge, the association between these two measures has been explored for the first time. This finding indicates that the two tests capture overlapping dimensions of navigational profiles and attitudes, but not so overlapping as to be equivalent.

We found that those that used the landmark strategy on RAM levels performed significantly better on both RAM and wayfinding levels than those who used the counting strategy. This mirrors recent evidence from the large participant group of over 37,000 participants (West et al., 2023). By combining the NSQ with the RAM tests for this study we reveal, for this population, that landmark-counting strategies appear to be orthogonal to the survey-route dimension captured by the NSQ. Notably, the lack of a correlation between wayfinding inefficiency and the RAM measures in this small lab sample matches our recent report of an absence of correlation between these in an online sample of over 37,000 participants (West et al., 2023). The absence of a correlation of the PI measure and the other navigation tasks is notable in light of recent evidence that PI, rather than other spatial tests, might be particularly important for early detection of AD (Kunz et al., 2015; Bierbrauer et al, 2020; Newton et al., 2023).

There are a number of limitations to our study that should be considered. Firstly, our sample size was limited by the challenge of conducting in lab testing, which was central to our planned design. Future studies with large online samples could explore a broader range of ages, and compare gender and performance across nations (Spiers et al., 2023). Secondly, the visuo-spatial tasks we used here were adapted from clinically used tests aimed at detecting differences between groups, whereas our aim was to explore variation in the population. Thus, it may be useful to further develop such tests to help optimally explore individual differences (Newcombe, Hegarty & Uttal, 2023). Finally, as with all correlative approaches to assessing individual differences, the capacity of the tests used will depend partly on the variance with the data generated by the test in the group tested. In our current study, the PI task had less variability than others and thus this may have impacted our capacity to detect some relationship between PI and other measures. Nonetheless, we were able to observe predicted correlation between this test and the D-Corsi, indicating the variation present was still sufficient to detect some effects.

In conclusion, our study highlights that many tests and self-rating scales for navigation and visuospatial abilities can be highly non-overlapping, at least in a UK university student sample. Our results help further characterise what the different tests used in SHQ are related to. For example, we show that wayfinding has some overlap with object-based visuospatial tasks, whereas PI and RAM tasks have much less. In future research it would be useful to probe a broader range of environments to understand how the complexity of the layouts (Coutrot et al., 2022a), and landmark density (Yesiltepe et al., 2019, 2021), lead to higher or lower correlations with object-based visuospatial ability. It would also be interesting to consider twin study cohorts where the genetic contribution to performance can be explored, and also expand to incorporate a measure of general cognitive ability (Malanchini et al., 2021). Finally, going beyond scores of task performance to modelling the behaviour using reinforcement learning methods (de Cothi et al., 2022; Lancia et al., 2023) holds much promise for helping obtain a more mechanistic understanding of the processes involved in navigation and visuo-spatial cognition.

## Acknowledgements

We would like to extend a thank you for all the participants who volunteered in this research.

## Appendix

## Gender-related Analysis

Though this paper was not specifically designed to measure gender differences, we explored gender-specific responses as part of a thorough investigation of the dataset. Gender-related analysis revealed significant differences in performances with respect to SHQ tutorial duration, RAM errors, NSQ score and SHQ stress rating (see Appendix Table 1). Male participants made significantly fewer RM and SWM errors on RAM levels than female participants. They were also significantly less stressed, and spent less time learning the controls of the game. Further, male participants had a significantly stronger mapping tendency as demonstrated by the NSQ. There was no association between gender and the type of strategy used to navigate RAM levels (*p* > .05).

**Appendix Table 1.** Metrics found to be significantly different for male participants (n = 20) versus female participants (n = 58) across measures.

| Measure | Female |  | Male |  | Statistics |  |  |  |
| --- | --- | --- | --- | --- | --- | --- | --- | --- |
|  | *M* | *SD* | *M* | *SD* | *t* | *df* | *p* | *d* |
| RAM RM Errors | 5.34 | 2.51 | 4.07 | 2.61 | 1.91 | 75 | .029 | .499 |
| RAM SWM Errors | 1.07 | 1.18 | .57 | .84 | 1.71 | 75 | .045 | .445 |
| SHQ Tutorial Duration | 47.36 | 6.81 | 44.07 | 4.61 | 1.94 | 72 | .028 | .519 |
| NSQ Score | -2.44 | 4.80 | -.10 | 5.37 | -1.81 | 75 | .037 | -.472 |
| SHQ Stress Rating | 4.44 | 2.39 | 2.60 | 1.50 | 3.21 | 75 | <.001 | .836 |

## Wayfinding and Path Integration Performance

We investigated the correlation between wayfinding and PI performance in the large Sea Hero Quest dataset (Coutrot et al., 2022a). We wanted to quantify how this correlation varies across training, wayfinding and path integration levels of different difficulties. We selected the participants who completed all Sea Hero Quest levels and were between 19 and 70 years old (N = 12,111, 5,979 males, mean age = 40.08, SD = 15.15 years). First, we computed the correlation between the overall PI performance (sum of correct answers across all PI levels) and the trajectory length of each WF level (see *Appendix Figure 1*). As expected, we found a small correlation for training levels 1 and 2 (*r* = -0.07) and a stable moderate correlation for all wayfinding levels (*r* ∼ -0.3). Then, we computed the point-biserial correlation between the overall training and wayfinding performance (the 1st component of a PCA across the trajectory lengths of the training levels on one side, and of the wayfinding levels on the other side) and the Boolean vector of correct answers for each PI levels (see *Appendix Figure 2*). As expected, we found a small correlation between training and PI performance and a stable moderate correlation between wayfinding and PI performance for the first 10 PI levels (until level 49). For the next PI levels, the magnitude of the correlation unexpectedly decreases to reach the same level as with the training performance.

**Appendix Figure 1.**
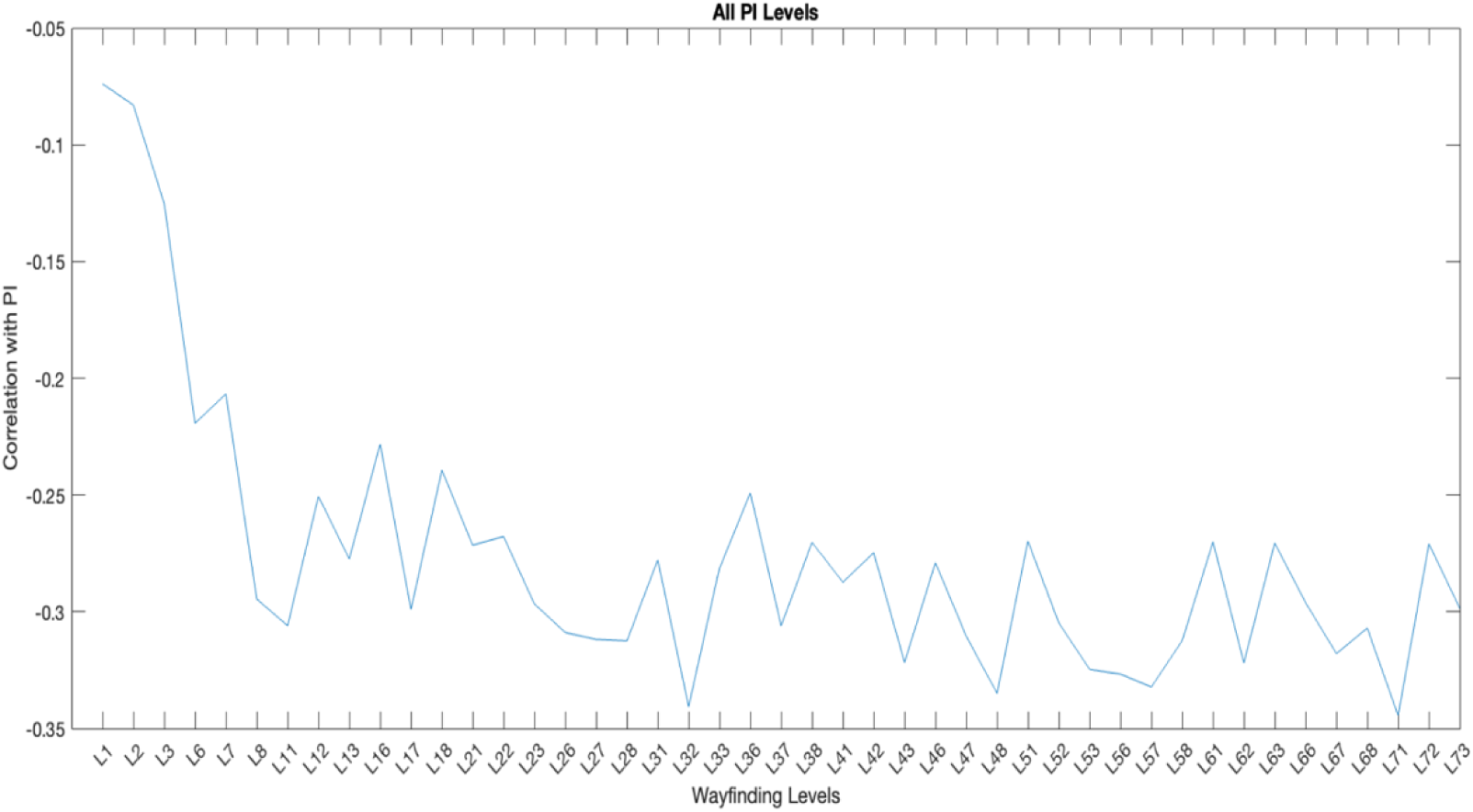
Correlation between the overall Path Integration performance and the trajectory lengths of each Wayfinding level.

**Appendix Figure 2.**
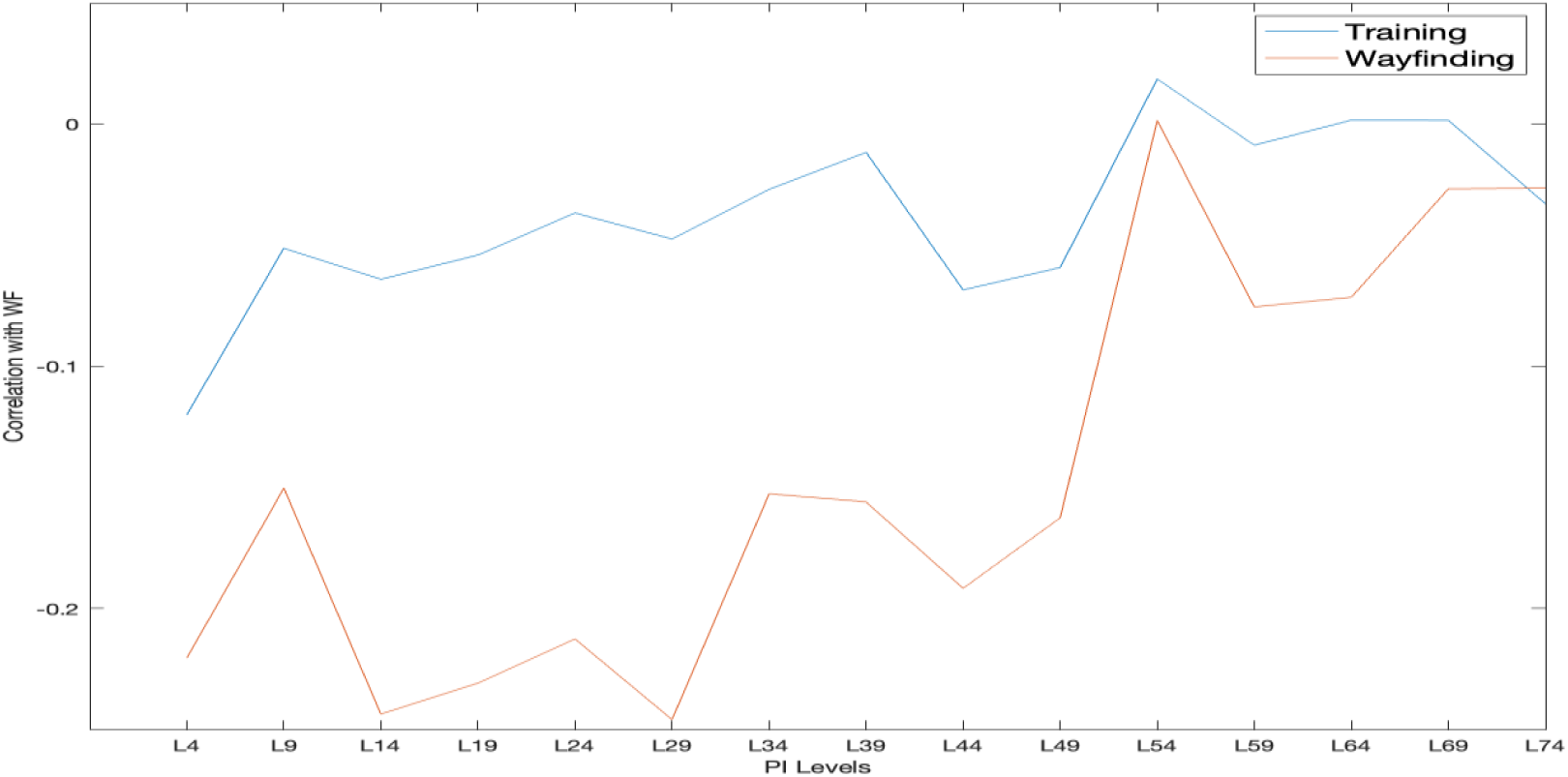
Point-biserial correlation between the overall Training and Wayfinding performance and the Boolean answers for each Path Integration level.

